# An M13 phagemid toolbox for engineering tuneable DNA communication in bacterial consortia

**DOI:** 10.1101/2025.06.22.660937

**Authors:** Abhinav Pujar, Anchita Sharma, Hadi Jbara, Manish Kushwaha

## Abstract

Intercellular communication is essential for distributed genetic circuits operating across cells in multicellular consortia. While diverse signalling molecules have been employed—ranging from quorum sensing signals, secondary metabolites, and pheromones to peptides, and nucleic acids—phage-packaged DNA offers a highly programmable method for communicating information between cells. Here, we present a library of five M13 phagemid variants with distinct replication origins, including those based on the Standard European Vector Architecture (SEVA) family, designed to tune the growth and secretion dynamics of sender strains. We systematically characterize how intracellular phagemid copy number varies with cellular growth physiology and how this, in turn, affects phage secretion rates. In co-cultures, these dynamics influence resource competition and modulate communication outcomes between sender and receiver cells. Leveraging the intercellular CRISPR interference (i-CRISPRi) system, we quantify phagemid transfer frequencies and identify rapid-transfer variants that enable efficient, low-burden communication. The phagemid toolbox developed here expands the repertoire of available phagemids for DNA-payload delivery applications and for implementing intercellular communication in multicellular circuits.

## Introduction

The rapid advancement of synthetic biology has enabled the construction of increasingly sophisticated genetic circuits capable of processing environmental information and responding in a programmable manner. These advances have resulted in applications such as detection of pollutants and disease biomarkers [1,2], smart therapeutics [3], and information processing logic circuits [4–6]. Early successes focussed largely on unicellular systems, where the complexity of circuits is inherently constrained by metabolic burden, limited availability of orthogonal genetic parts, and undesirable crosstalk between circuit components. More recently, synthetic genetic circuits have begun to move towards multicellular architectures [7–11]. In this distributed multicellular computing paradigm, circuit sub-functions are executed by different strains within a synthetic consortium. Distributing circuit elements across a population of interacting cells offers several advantages, including reduced metabolic burden per cell, minimisation of component crosstalk, and enhanced modularity, while also opening up opportunities for specialisation, concurrency, and fault tolerance [12,13]. Such multicellular consortia allow for scalable and parallelizable designs; however, the success of these distributed circuits critically depends on reliable and programmable intercellular communication to coordinate behaviour across the circuit’s constituent parts.

Intercellular communication—analogous to external “wiring” in electronic circuits—links the inputs and outputs of sub-circuits distributed across different cells. Without such communication, coordination between cell types and collective decision-making would not be possible. To enable this, synthetic biologists have repurposed a wide range of natural signalling molecules. Quorum sensing systems, particularly those based on bacterial homoserine lactones (HSLs) [7,14–17], have been the most commonly used. Other communication strategies have included yeast mating pheromones [8], mammalian receptor-ligand pairs [9], small secondary metabolites [18], and secreted peptide signals [19]. However, each of these systems is typically capable of transmitting only a single signal for each type of communication channel [20], and the repertoire of available orthogonal channels is small—with typically fewer than ten distinct, non-crosstalking signals. As the scale and sophistication of multicellular circuits increase, these limitations in signal orthogonality and bandwidth become critical bottlenecks.

To overcome the constraints of small-molecule signalling, researchers have looked to nucleic acids—specifically DNA—as signals for intercellular communication [20]. In nature, DNA is frequently exchanged between microbes through horizontal gene transfer mechanisms such as transformation, conjugation, and phage-mediated transduction [21]. These processes can be used to modify undomesticated bacteria [22], for phage therapy [23], inactivation of antimicrobial resistance [24], and metabolic pathway delivery [25]. Importantly, DNA can encode highly complex, information-rich messages whose expression can alter the functional state of recipient cells. DNA-based communication offers several advantages: message content can be easily programmed; orthogonality can be introduced through sequence modification; and a single communication channel (e.g., phage transduction) can transmit a wide variety of messages. Several recent studies have demonstrated the feasibility of DNA-based messaging for synthetic circuits. Conjugation-based systems have been used to target pathogens [26] and targeted DNA delivery to specific cells within a population [27]. Meanwhile, the filamentous bacteriophage M13, which continuously packages and secretes DNA-containing phage particles without lysing its host [28], has been used to implement phage-mediated messaging and distributed circuits. M13 can package plasmids known as phagemids, which contain an M13 packaging signal and any additional desired genetic content. This approach was used in one of the earliest demonstrations of DNA-based intercellular communication, where a phagemid carrying a T7RNAP gene was transferred to a receiver cell, triggering gene expression [20]. More recent works have demonstrated multicellular logic circuits using phage-mediated intercellular CRISPRi for biocomputation in bacterial consortia [29,30].

Despite its promise, broader adoption of M13-based DNA messaging is limited by practical challenges. Most notably, implementing multiple parallel message transfers to a single receiver cell requires the stable coexistence of multiple phagemids. This is difficult because plasmids that share the same replication origin are incompatible: they compete for the same replication machinery, leading to instability or loss [31]. To address this limitation, here we developed a modular phagemid toolbox comprising five variants, each harbouring a distinct replication origin. These origins are maintained at different plasmid copy numbers and impose varying physiological burdens on the host, which in turn affect the rate of phagemid secretion and the efficiency of intercellular communication. By characterizing these differences, we provide circuit designers with a tuneable, orthogonal set of DNA-based communication signals. Using an intercellular CRISPR interference (i-CRISPRi) system, we demonstrate that these phagemid variants exhibit secretion rates that vary over ∼3000-fold per cell and transfer frequencies that vary over 180-fold. Our results reveal that the choice of replication origin not only affects intracellular plasmid maintenance but also significantly influences intercellular communication dynamics. This expanded phagemid library provides a flexible and modular platform for encoding and communicating multiple, distinct messages in multicellular circuits, advancing the design of scalable, programmable, and information-rich synthetic systems.

## Results

### Construction and characterisation of an M13 phagemid toolbox

The naturally occurring filamentous bacteriophage M13 infects *F*+ *E. coli* cells by binding to the F-pilus as its primary receptor. Upon entry, the single-stranded phage DNA (ssDNA) is converted into a double-stranded replicative form (RF), which is amplified by the host’s replication machinery. The phage DNA is subsequently mobilised for packaging via a specific M13 packaging signal (ps) [32,33]. M13 phagemids, first engineered in 1985 [34], combine a bacterial origin of replication with the phage DNA packaging signal, enabling stable maintenance in the bacterial host as well as their packaging into secreted phage particles, provided helper machinery is also provided in the same cell. Phagemids have been widely used for applications in nanotechnology [35], phage display [36], vaccine development [37], biosensing [38] and directed evolution [39].

In previous work [40], we performed a detailed characterisation of secretion and infection rates of an M13 phagemid containing the pUC replication origin, based on pSB1K3 [41]. In this work, we have constructed four new phagemid variants with replication origins pBR322, p15A, pColA, and pSC101, based on the Standard European Vector Architecture (SEVA) vectors pSEVA291, pSEVA261, pSEVA29A1, and pSEVA271, respectively [42]. Differing only in the bacterial origin of replication, the five phagemid variants were constructed to contain the M13 packaging signal, and coding sequences for *kanR*, *lacZ*α, and *gp3* genes (Methods, Fig 2a). The minor coat protein gene *gp3* was included on the phagemids to enable more accurate counting of phage titres by plaque-forming unit (PFU) assays, since phagemids lacking *gp3* can only be assayed by the less accurate colony-forming unit (CFU) assays [40]. The resulting phagemids were called +gp3_pUCφ,+gp3_pBR322φ, +gp3_p15Aφ, +gp3_pColAφ, and +gp3_pSC101φ (Supplementary Info.). Next, this toolbox of phagemid variants was characterised for its growth, sender cell phagemid copy number, phage secretion dynamics, intercellular communication, and transfer frequencies (Fig. 1).

**Figure 1.**
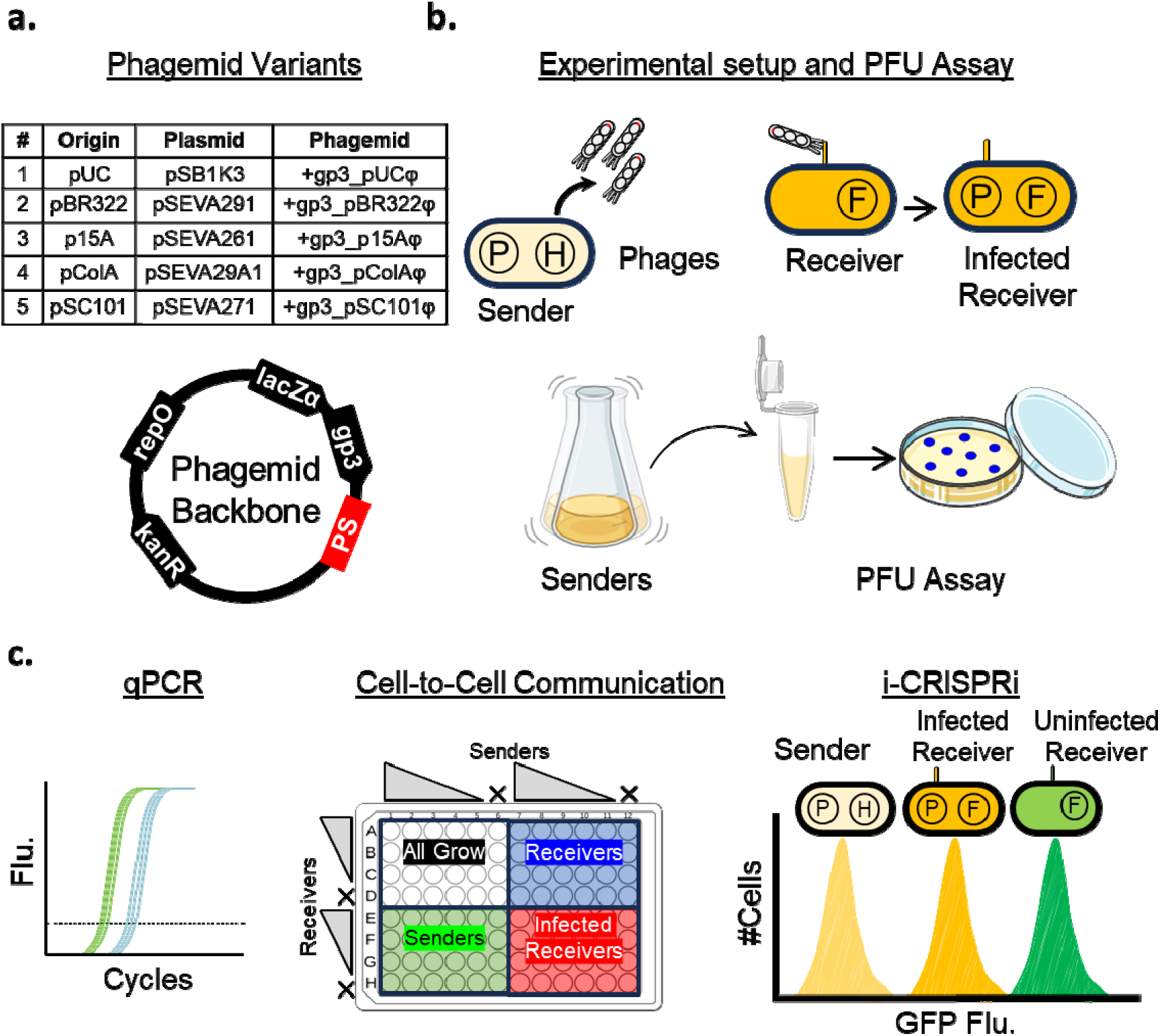
Schematic of phagemid toolbox variants and the experimental setup used for their characterisation. **(a)** Phagemids used in this study carry one of five different replication origins, an M13 packaging signal (PS), and several other genes: *lacZ*_α_, *gp3*, *kanR*. **(b)** Sender variants were evaluated for growth by periodically measuring OD_600_ and measuring phage concentrations using the PFU Assay. **(c)** Intracellular phagemid copy numbers were determined by qPCR. Sender and receiver cells were co-incubated to evaluate growth and resource competition (plate reader), as well as transfer frequencies (flow cytometry) by intercellular CRISPRi (i-CRISPRi). Some images used in this figure were taken from <u>bioicons.com</u> and are licensed under CC-BY 3.0, CC0, and MIT.

**Figure 2.**
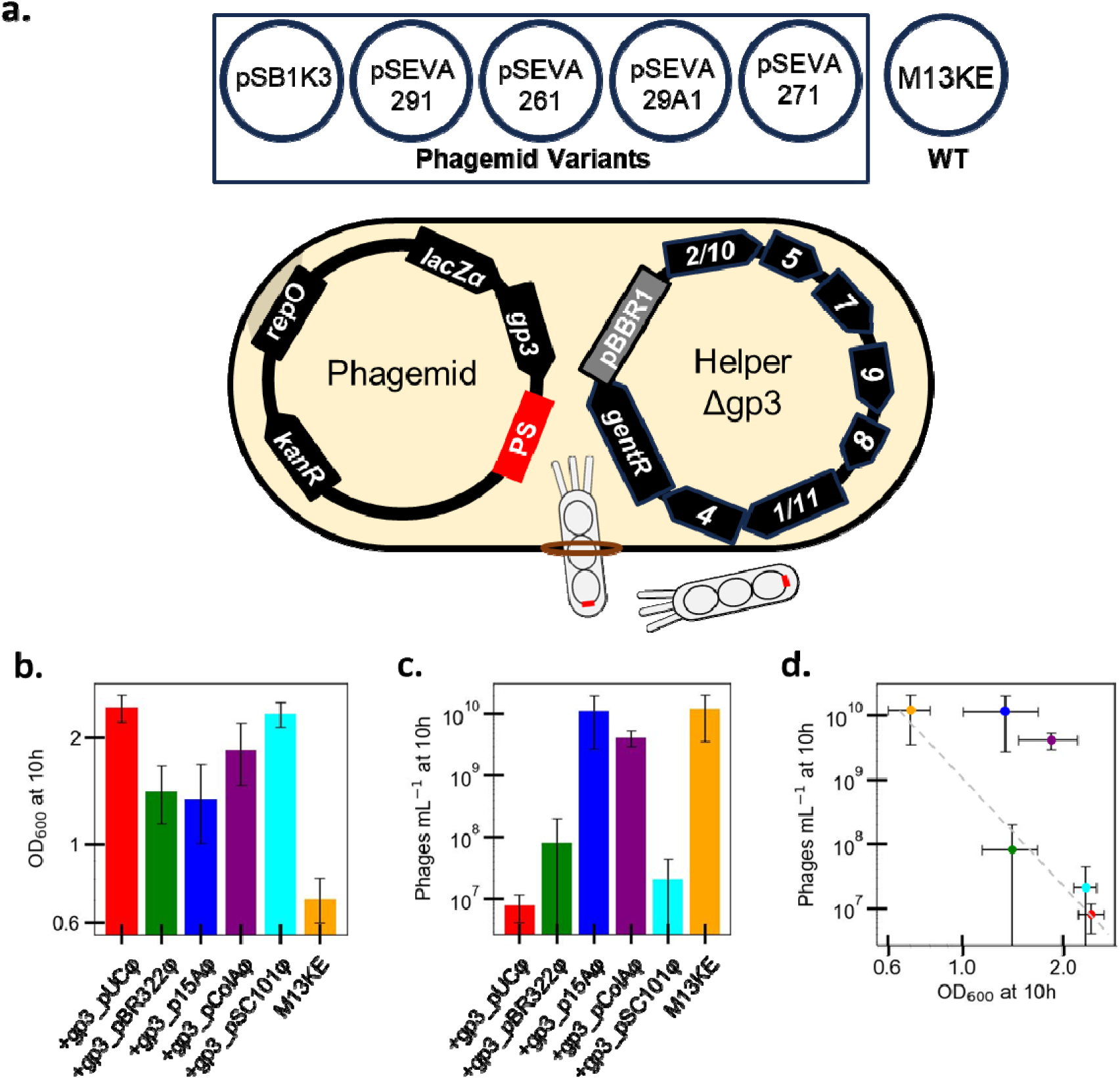
Culture densities and phage yields of sender variants in the phagemid toolbox. **(a)** Sender cell consists of two plasmids: the helper plasmid (pSEVA63_HelperΔgIII [40]) and the phagemid. **(b-d)** Sender cells with different phagemids were grown in selection media for ∼10 hours. Final cell densities (OD_600_) and phage yields are plotted in (b) & (c), respectively. The two are plotted against each other in (d).

In previous work [40], we performed a detailed characterisation of secretion and infection rates of an M13 phagemid containing the pUC replication origin, based on pSB1K3 [41]. In this work, we have constructed four new phagemid variants with replication origins pBR322, p15A, pColA, and pSC101, based on the Standard European Vector Architecture (SEVA) vectors pSEVA291, pSEVA261, pSEVA29A1, and pSEVA271, respectively [42]. Differing only in the bacterial origin of replication, the five phagemid variants were constructed to contain the M13 packaging signal, and coding sequences for *kanR*, *lacZ*α, and *gp3* genes (Methods, Fig 2a). The minor coat protein gene *gp3* was included on the phagemids to enable more accurate counting of phage titres by plaque-forming unit (PFU) assays, since phagemids lacking *gp3* can only be assayed by the less accurate colony-forming unit (CFU) assays [40]. The resulting phagemids were called +gp3_pUCφ, +gp3_pBR322φ, +gp3_p15Aφ, +gp3_pColAφ, and +gp3_pSC101φ (Supplementary Info.). Next, this toolbox of phagemid variants was characterised for its growth, sender cell phagemid copy number, phage secretion dynamics, intercellular communication, and transfer frequencies (Fig. 1).

Sender cells were constructed by co-transforming *E. coli* Top10 cells with a helper plasmid (pSEVA63_HelperΔgIII [40]) and one of the five phagemid variants (Fig 2a). Senders, including a full phage M13KE control, were cultured in LB media with appropriate antibiotics for ∼10 hours. The growth densities achieved by the different sender variants vary over 3.48-fold (Fig. 2b). The highest densities are observed in the sender variants +gp3_pUCφ and +gp3_pSC101φ, with OD_600_ values of 2.44 and 2.34, respectively, followed by +gp3_pColAφ at 1.84, +gp3_pBR322φ at 1.41, and +gp3_p15Aφ at 1.35. The lowest density was observed for the full phage M13KE at 0.7. The phage yield from the different sender cultures also shows a marked variation of >500-fold at the end of 10 hours (Fig. 2c). Sender variants +gp3_p15Aφ, M13KE, and +gp3_pColAφ achieve the highest phage titers, with yields of 1.14 × 10^10^, 1.20 × 10^10^, and 4.15 × 10^9^ PFU mL^-1^, respectively. In comparison, the +gp3_pBR322φ variant produces a moderate yield of 8.08 × 10^7^ PFU mL^-1^, while the +gp3_pSC101φ and +gp3_pUCφ variants secrete the lowest phage titers at 2.10 × 10^7^ and 7.93 × 10^6^ PFU mL□¹, respectively. These data show that there is an inverse relationship between phage yields and end-point cell densities reached by the phage-producing batch cultures. Two phagemid variants (+gp3_p15Aφ and +gp3_pColAφ) seem to be exceptions, with high yields comparable to phage M13KE but also higher final ODs, suggesting that phagemid replication origins p15A and pColA can help achieve both high phage titres and high sender cell densities.

### Intracellular phagemid copy numbers impact phage secretion rates

To measure growth rates, cultures of sender variants were grown in LB media with appropriate antibiotics for 10 hours, while a spectrophotometer (OD_600_) reading was recorded at regular intervals (Fig 3a). At each time-point, phages were isolated from a 1 mL culture sample and the bacterial pellet frozen (Methods). Growth rates were calculated from the OD_600_ data, both for the log and the early stationary phases (Fig 3b). PFU assay was performed using the isolated phage samples to obtain secretion curves (Fig 3c), and calculate per cell secretion rates in the log and the stationary phases (Fig 3d). The data obtained are summarised in Table 1. We found that phage secretion rates peak during the exponential (log) growth phase (Fig. 3a,d). The secretion rates of sender variants +gp3_p15Aφ and +gp3_pColAφ are 8.15 × 10^-1^ and 1.57 × 10^-1^ cell^-1^ min^-1^, respectively, significantly higher than those of lower-yielding variants such as +gp3_pSC101φ and +gp3_pUCφ, which exhibit rates of 1.54 × 10^-3^ and 2.77 × 10^-4^ cell^-1^ min^-1^, respectively (Fig. 3d). In the stationary phase, when cellular growth slows down, secretion rates drop significantly across all sender variants. For example, +gp3_p15Aφ and +gp3_pColAφ have secretion rates of 5.99 × 10^-2^ and 4.45 × 10^-2^ cell^-1^ min^-1^, respectively, which are lower than their corresponding rates during the log phase (Fig. 3d).

**Figure 3.**
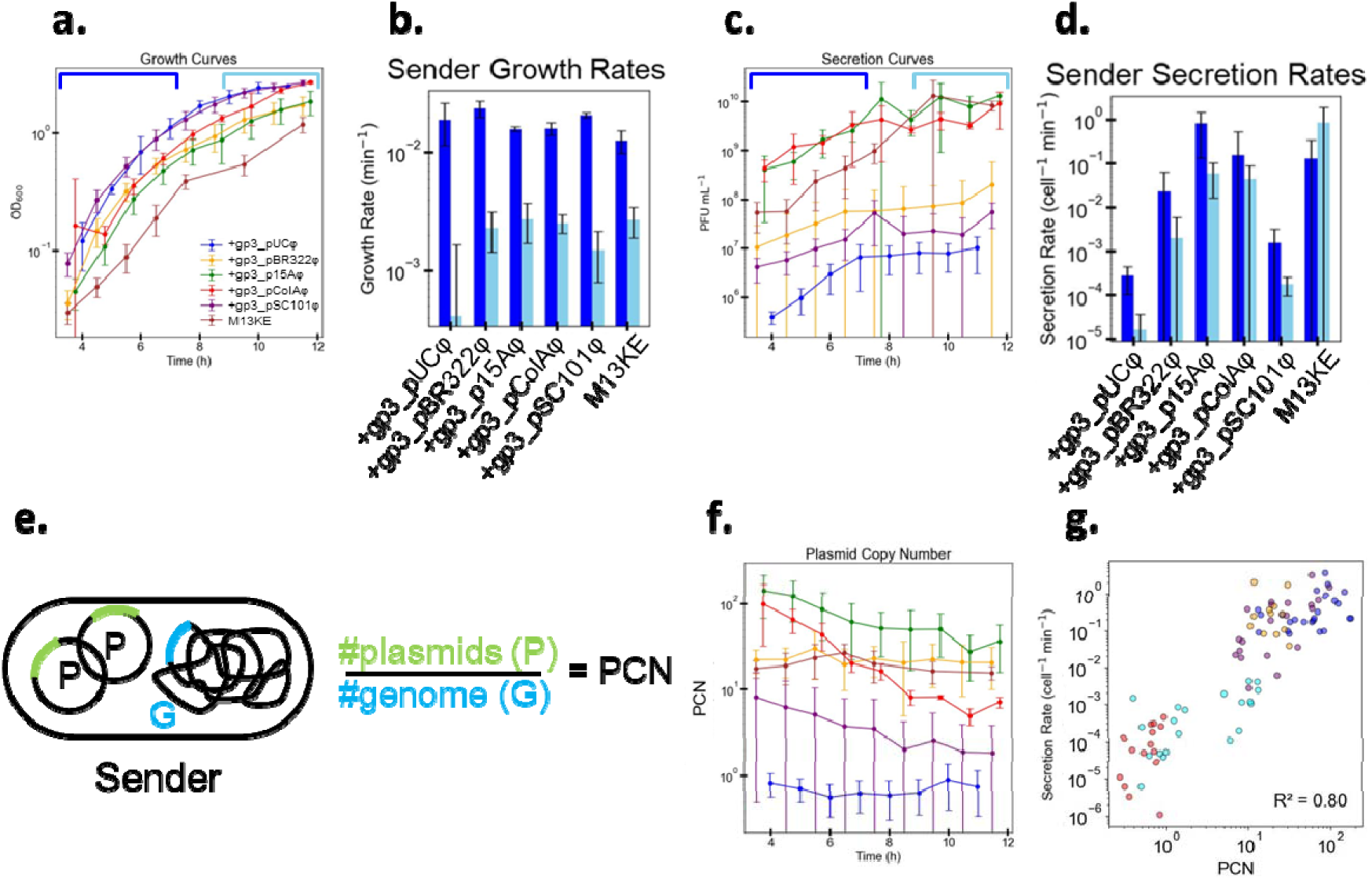
Time-course analysis of the phagemid toolbox senders. **(a)** Growth curves were obtained by recording OD_600_ at intervals of 1 hour. Log phase region is marked in dark blue, while the stationary phase region is marked in light blue. **(b)** Instantaneous growth rates of all sender cultures in the log and the stationary phases of growth. **(c)** Secretion curves were obtained by performing the PFU assay on phage isolates collected at intervals of 1 hour. **(d)** Secretion curves (c) and instantaneous growth rates (a) were used to calculate the instantaneous secretion rates plotted here for the log and the stationary phases of growth. **(e)** Quantitative PCR (qPCR) was performed on cells isolated from culture samples in (a). The plasmid copy number (PCN) was determined as the ratio of plasmid DNA concentration and genomic DNA concentration in the same sample. **(f)** PCNs were estimated for each phagemid variant using cells isolated from culture samples in (a) at several time-points. **(g)** PCN and instantaneous secretion rates at a given time-point were plotted against each other for all sender variants at all time-points, except +gp3_pBR322_φ_.

**Table 1.**
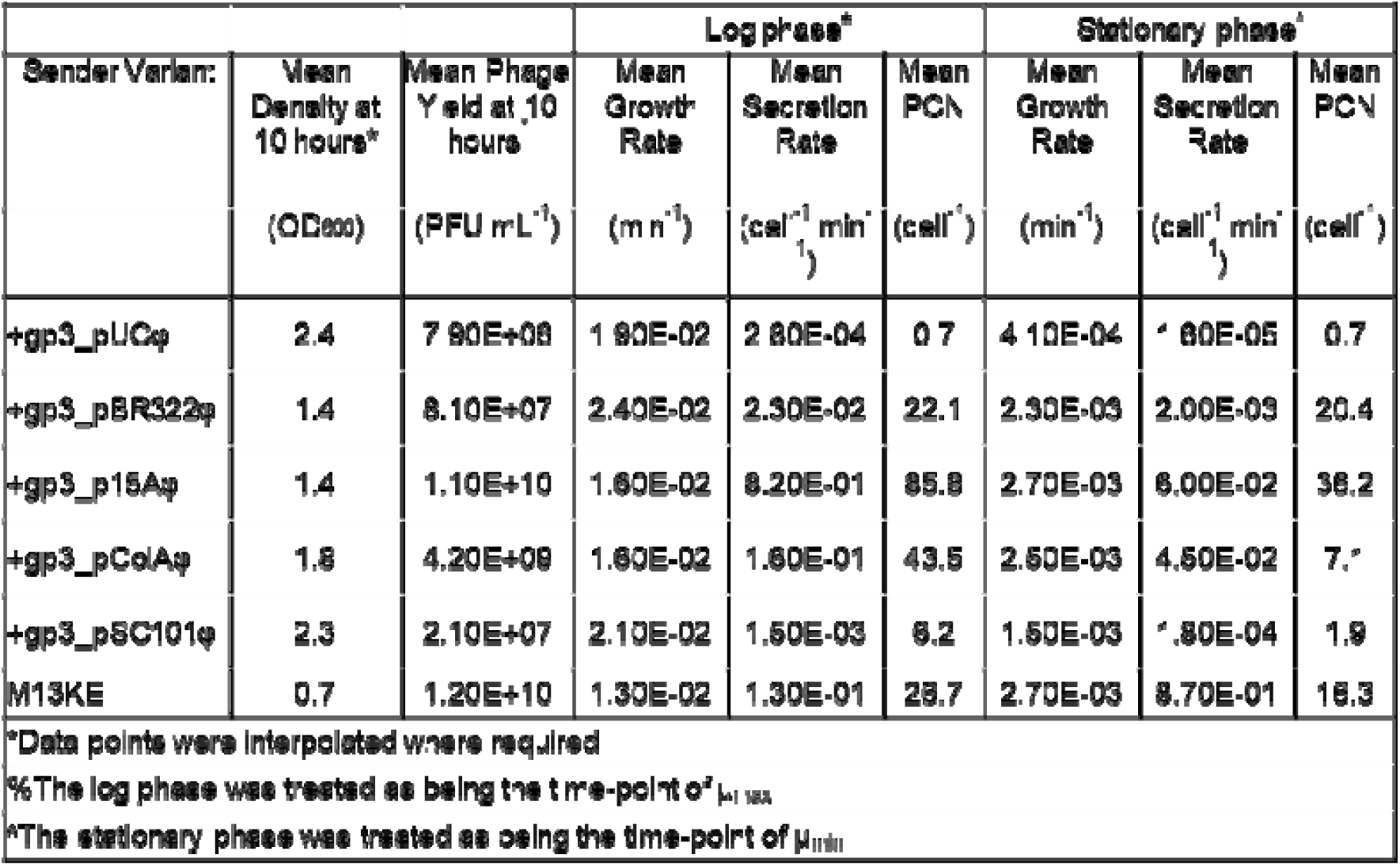
Kay characaratlct of sardar variants darlvad from growth one sacratlon dots.

The previously frozen sender cell pellets (Fig 3e) were thawed and used for plasmid copy number (PCN) determination using qPCR (Methods). Values were calculated as the number of phagemid copies per genome in the sender cells, and plotted for each time-point (Fig 3f). During the log phase, the highest-producing sender variants +gp3_p15Aφ and +gp3_pColAφ maintain 85.84 and 43.54 phagemid copies per cell, respectively, whereas the lowest-producing sender variants +gp3_pUCφ and +gp3_pSC101φ maintain only 0.70 and 6.16 copies per cell (Fig. 3f). For some sender variants, PCNs decrease significantly in the stationary phase. For instance, the PCNs for +gp3_p15Aφ and +gp3_pColAφ reduce to 36.23 and 7.05 in the stationary phase, respectively. In contrast, the PCNs for +gp3_pUCφ and +gp3_pBR322φ remain relatively constant, changing slightly from 0.70 to 0.74 and 22.14 to 20.4. This stability in PCN indicates that some replication origins maintain consistent phagemid copy numbers, even during transitions in cellular physiology.

Table 2 shows plasmid copy numbers reported in previous studies using ddPCR [43], fluorescence microscopy [44] or qPCR [45,46] alongside those determined in this work by qPCR. Most of the PCN estimations reported differ significantly across studies. This could be due to a variety of reasons such as plasmid size, cargo carried on them, growth medium, bacterial strains used and different methods used to quantify PCNs [43,44].

**Table 2:** Plasmid Copy Numbers of Different Replication Origins.

| **Table 2: Plasmid Copy Numbers of Different Replication Origins** |  |  |  |  |  |  |
| --- | --- | --- | --- | --- | --- | --- |
| Replication Origin | Jahn et al. 2016 | Shao et al. 2021 | Lee et al. 2006 | Song et al. 2017 | This Study (Log Phase) | This Study (Stationary) |
| p15A | 8.6 ± 1.9 | 9 ± 11 | — | 50 | 85.8 ± 48.5 | 36.2 ± 20.6 |
| pSC101 | 3.4 ± 0.5 | 4 ± 4 | — | — | 6.2 ± 6.4 | 1.9 ± 1.9 |
| pColA | — | — | — | 12 | 43.5 ± 15.5 | 7.1 ± 1.0 |
| pUC | 8.9 ± 4.5 | 61 ± 60 | — | — | 0.7 ± 0.2 | 0.7 ± 0.4 |
| pBR322 | — | — | 18.5 | 38 | 22.1 ± 8.5 | 20.4 ± 10.5 |
| M13KE | — | — | — | — | 26.7 ± 7.1 | 16.3 ± 4.8 |

Secretion rates, growth rates, and plasmid copy number (PCN) of all sender variants were plotted against each other to identify their relationships (Supplementary Info.). PCN and secretion rates in the log-phase show a strong positive relationship (Supplementary Info.), but other variables did not show any clear global trends. This suggests that secretion rates vary with changing growth physiology due to associated changes in the intracellular copy number of phagemids. As the intracellular DNA substrate available for packaging increases, so does the phage secretion rate. The PCN-secretion rate relationship appears even stronger (increase in R^2^ from 0.55 to 0.80; Supplementary Info.) when the outlier +gp3_pBR322φ is excluded from the analysis (Fig. 3g). This suggests that additional factors other than PCN play a role in the packaging and secretion of the pBR322 phagemid.

### Phagemid variants exhibit different resource competition and intercellular communication dynamics

Having found above that per cell phage secretion rates vary over ∼3000-fold depending on the phagemid variant, we next investigated how these differences influence intercellular communication dynamics between phage-secreting sender cells and susceptible receiver cells growing together in a co-culture. Senders and receivers were co-incubated without antibiotics for 1 h, followed by growth under four conditions: without antibiotics, or with antibiotic/s that select for receivers only (tetracycline), senders only (gentamycin) or infected-receivers only (kanamycin + tetracycline) (Fig 4a). Growth was monitored using optical density (OD_600_) measurements taken periodically for over 18 hours (Supplementary Info.).

**Figure 4.**
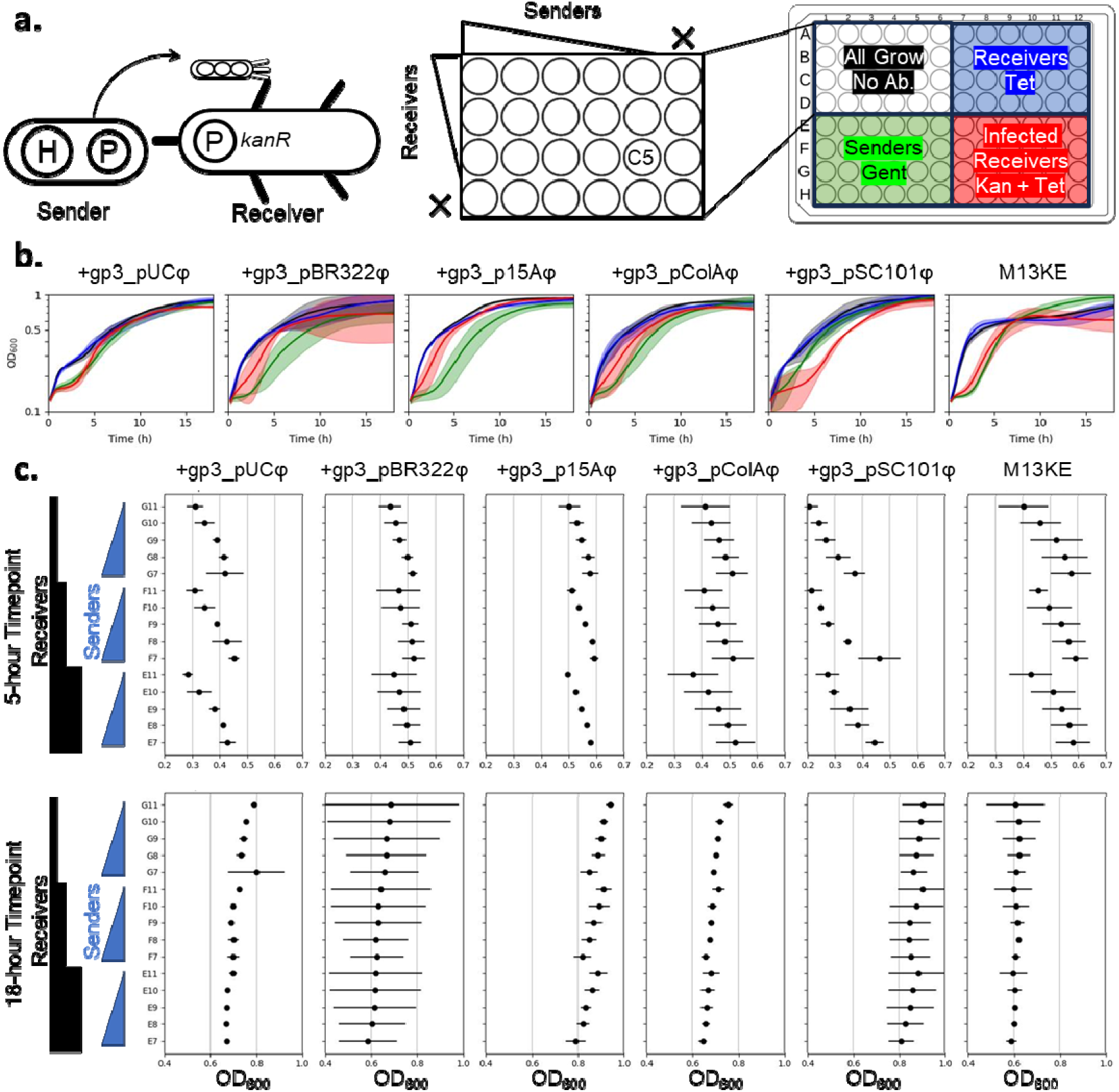
Cell-to-cell communication dynamics of different sender variants between growing sender and receiver cells. **(a)** Sender and receiver cells with different starting densities were co-cultured in a plate reader for 1 hour without selection, and then antibiotic selection was applied to select receivers only (tetracycline), senders only (gentamycin), infected receivers only (kanamycin+tetracycline). A no-antibiotic control was also taken along. **(b)** Growth curves obtained in each selection condition were plotted for the well that started with the lowest cell densities (well C5). Growth curves in blue represent receiver-only growth, green represents sender-only growth, red represents infected receiver-only growth, and black represents no selection. **(c)** Plots of cell densities (OD_600_) reached at a middle (5 h, top panel) and the end (18 h, bottom panel) time-points in wells with different starting sender and receiver densities for all phagemid variants in the infected receivers-only condition. X-axis shows the density reached, and y-axis shows the starting sender and receiver cell densities added. The +gp3_pUC_φ_ (pSB1K3) data presented for comparison here is from [40].

Growth dynamics in these conditions revealed how resource competition and antibiotic selection impacted intercellular communication (Fig 4b, Supplementary Info.). As seen in our previous work [40], in the absence of antibiotic selection both sender and receiver co-cultures exhibited monophasic growth kinetics. Similarly, cultures in receiver-only (tetracycline) condition show monophasic growth, suggesting that sender cells are rapidly eliminated by tetracycline treatment, with subsequent growth driven exclusively by the receiver population. In co-cultures where two cell types compete for shared resources, biphasic growth curves typically indicate sequential population dynamics, with one cell type initially dominating followed by the delayed growth of the other [47]. Such biphasic growth pattern was evident in the senders-only (gentamycin) condition, particularly at low initial sender and receiver densities (Figure 4b). The delayed killing of receiver cells by gentamycin allows them to transiently grow and compete for resources before their eventual death.

Biphasic growth kinetics is also seen in infected-receivers-only (kanamycin + tetracycline) condition (Fig 4b), where only receiver cells infected during the 1 h preincubation grow after the kanamycin + tetracycline selection is applied. To further understand the contributions of sender and receiver inoculation numbers to the communication and selection process, we plotted ODs of several wells at an early time-point (5 h) and the end-point (18 h) of post-selection growth (Fig 4c, Supplementary Info.). Sender variants +gp3_p15Aφ, +gp3_pColAφ, and M13KE with high secretion rates (Table 1) are expected to secrete high titres of phages that infect a higher number of receiver cells in 1 hour. In contrast, the +gp3_pUCφ sender with low secretion rates is expected to infect fewer receiver cells. Sender variants +gp3_pBR322φ and +gp3_pSC101φ with medium secretion rates are expected to produce moderate levels of phages that infect a moderate number of receiver cells. This is reflected in Fig 4c, where by 5 hours receivers incubated with sender variants +gp3_p15Aφ and M13KE reach the higher ODs of ∼0.58, followed by +gp3_pColAφ at ∼0.52, +gp3_pBR322φ at ∼0.49, +gp3_pSC101φ at ∼0.44, and +gp3_pUCφ at the lowest at 0.42. For all phagemids, a clear sender dose dependence was seen at the 5 h time-point: higher initial sender OD resulted in more infected receivers.

At 18 hours, the highest densities are achieved by receiver cells infected by phage variants +gp3_p15Aφ and +gp3_pSC101φ at ∼0.79, followed by +gp3_pUCφ at ∼0.67, +gp3_pColAφ at ∼0.64. The lowest densities were recorded for receivers infected by M13KE and +gp3_pBR322φ phages at ∼0.59. For most phagemid variants, the 18 h OD reaches similar values irrespective of the initial senders or receivers inoculated. However, for the high-secretion rate variants (+gp3_p15Aφ and +gp3_pColAφ) there is an inverse relationship between initial sender OD and final infected-receiver OD, indicating that early growth by these senders sufficiently depletes resources to prevent infected receivers from reaching higher final densities. Additionally, co-cultures with higher initial infection do not necessarily have higher final density.

### Diverse phagemid transfer frequencies estimated by intercellular CRISPRi

CRISPR interference (CRISPRi) has been used widely in synthetic biology to regulate gene expression in a wide range of organisms. CRISPRi system consists of a catalytically dead variant of the Cas9 nuclease (dCas9) and a single guide RNA (sgRNA) [48]. The dCas9 and sgRNA form a ribonucleoprotein complex that can bind to a 20 nt DNA sequence with high specificity. The target binding site directly depends on the programmable sgRNA sequence. When the binding site is inside or near a promoter, the dCas9-sgRNA complex prevents transcription from the promoter by blocking RNA polymerase access. In previous work, we have developed an intercellular CRISPRi (i-CRISPRi) system that enables repression of a GFP gene in receiver cells upon transfer of an sgRNA-encoding phagemid from sender cells (Fig 5a,[40]). The system enabled us to track the phagemid transfer frequency from senders to receivers in growing co-cultures without applying any antibiotic selection, unlike in the previous results section.

**Figure 5.**
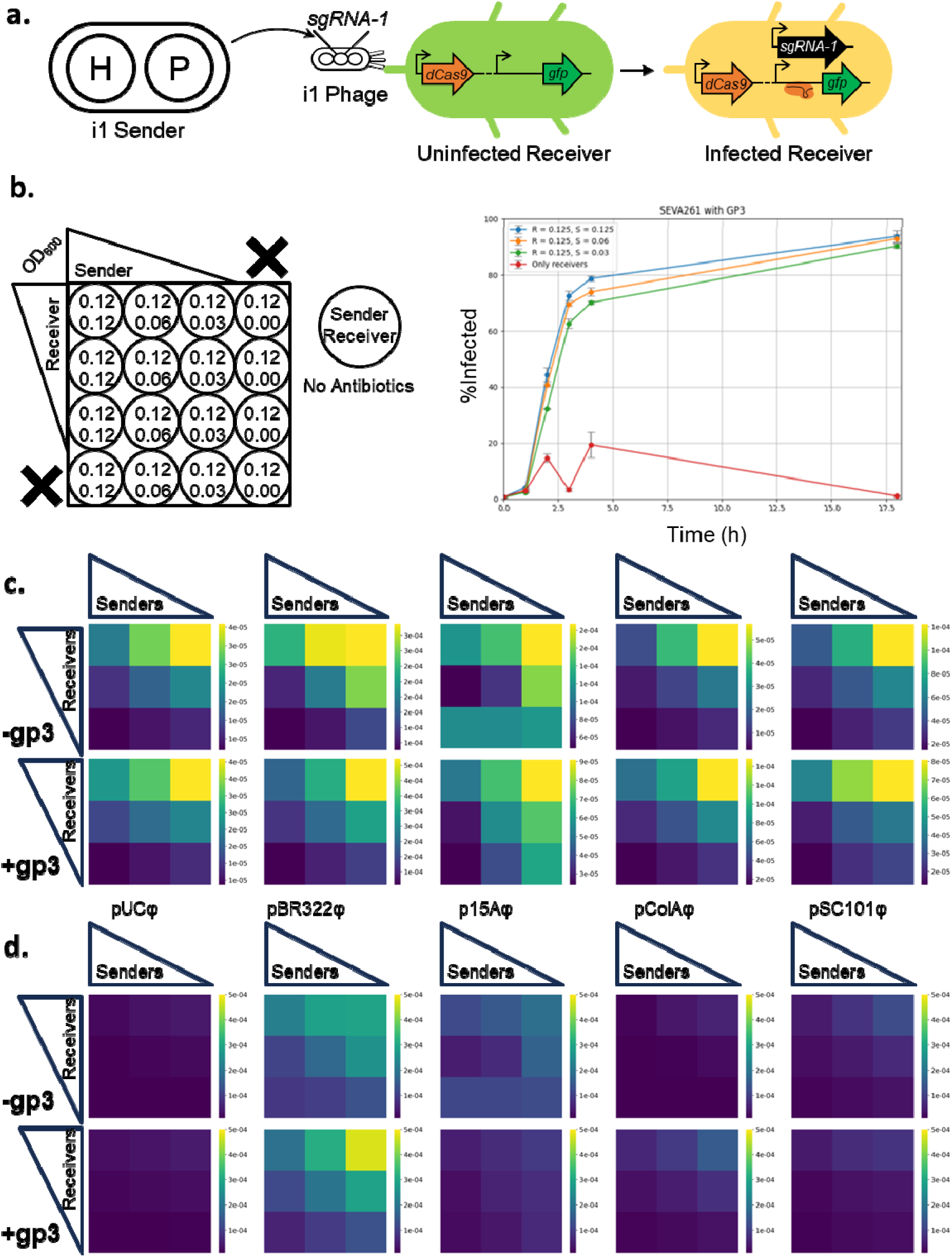
Intercellular CRISPRi using variants of the phagemid toolbox. **(a)** An sgRNA-1 sender library was created using all the toolbox phagemid variants with different replication origins, both in the-gp3 and +gp3 versions. Sender cells constitutively secrete phages that carry the *sgRNA-1* gene, and the F+ receiver cells constitutively express dCas9. Upon infection by the phage, the sgRNA is expressed in the receiver cell, which results in the repression of the previously expressing *gfp* gene. **(b)** Sender and receiver cells were grown together at different starting densities for 18 hours without antibiotics. At timepoints 0, 1, 2, 3, 4, and 18 hours, a sample of cell culture was used for flow cytometry. Using the flow cytometry data, we were able to differentiate between the number of infected and uninfected receiver cells. This was used to calculate the %Infected over time. **(c)** Transfer frequencies were calculated for all sender variants, and heatmaps for each set calculated separately. Only data from the wells that contain both senders and receivers is plotted. **(d)** Same data as in (c), with transfer frequencies for all sender variants with heatmaps re-scaled to a common scale.

In this work, sgRNA sequence (*sgRNA-1*) was cloned into all phagemid variants based on different vectors—pSB1K3, pSEVA291, pSEVA261, pSEVA29A1, and pSEVA271—in both the-gp3 and +gp3 versions. Sender variants were made by co-transforming *E. coli* Top10 cells with the phagemid and a helper plasmid. Receiver cells were F+ *E. coli* cells with the dCas9-GFP plasmid (TOP10F_dCas9-GFP_NOT_i1 [40]) that encodes for dCas9 expression from a constitutive promoter and GFP expression from a CRISPRi-repressible promoter. Senders and receivers in different starting ratios were co-incubated in a plate reader for 4 hours without antibiotics, and GFP expression monitored by flow cytometry from samples collected at 0, 1, 2, 3, and 4 hours (Fig. 5b, left). By appropriate gating of the flow cytometry data (Methods), the numbers of senders (no GFP), infected receivers (low GFP), and uninfected receivers (high GFP) could be determined.

In an example plot of the +gp3_p15Aφ phagemid data (Fig 5b, right), we found that for a fixed initial receiver OD of 0.125, increasing initial sender OD speeds up phagemid transfer slightly. However, for fixed initial sender OD of 0.125 and varying receiver ODs (Supplementary Info.), the differences are not apparent. This suggests that at even the lowest sender:receiver ratios, receiver cells get rapidly infected, particularly for high-secreting senders. These trends broadly hold for most phagemid variants (Supplementary Info.).

To better understand the role of inoculation ratios on phagemid-specific intercellular communication, we calculated transfer frequencies for all ratios using a method used to obtain transconjugation frequency in conjugation experiments [27]. The transfer frequencies obtained were plotted against time for all phagemids, and in both versions (-gp3, +gp3) (Supplementary Info.). We find that for phagemids p15Aφ, pSC101φ, pBR322φ, and pUCφ, transfer frequencies of the-gp3 version were higher than those of the +gp3 version. The exception was phagemid pColAφ, for which transfer frequencies of the +gp3 version were higher than those of the-gp3 version. These likely reflect the differences in expression levels of the essential gp3 gene in the sender cells, either from the phagemid (+gp3 version) or from the helper plasmid (-gp3 version).

Heatmaps of transfer frequencies at 3 h for all phagemid variants were plotted for comparison (Fig. 5c). Since there is no selection applied here, the data reflect only the growth dynamics of receivers and senders as they communicate with each other using phages. One would expect higher sender ODs to result in higher transfer, but that was not observed. Instead, the highest transfer frequencies were seen in the lowest-sender highest-receiver starting OD conditions. This suggests that when phage secretion rates are high, fewer senders can infect a sufficient number of receiver cells without causing substantial resource competition. These infected receivers then grow, resulting in vertical transmission of the phagemid and more infected receivers. When comparing transfer frequencies across all +gp3 phagemid variants (Fig. 5d), we see that +gp3_pBR322φ has the highest transfer frequency, which is interesting because this phagemid does not have high secretion rates (Fig 3d). Similarly, across all-gp3 phagemid variants (Fig. 5d), we also find that - gp3_pBR322φ has the highest transfer frequency.

## Discussion

In this work, we have built and characterised a modular phagemid toolbox for M13-based intercellular communication in *E. coli*, and compared it with the widely used full phage M13KE. By systematically varying the origin of replication while keeping the rest of the phagemid and the helper plasmid constant, we were able to interrogate how the plasmid copy number (PCN) in the sender cells impacts their growth rates, PCNs, the phage secretion rates, and in turn the intercellular communication efficiencies. Our time-resolved analysis of secretion rates and PCNs also revealed that not only is the relationship between copy number and secretion rate dynamic and growth phase-dependent, it is also replication origin specific. As growth transitions into the stationary phase, we observed significant reductions in both PCN and secretion rate. This suggests that while high-copy origins drive elevated phage output during active growth, their advantage diminishes as cells exit the log phase. Phagemids with high intracellular copy numbers such as +gp3_p15Aφ and +gp3_pColAφ led to the highest phage yields and secretion rates, with ∼3000-fold differences observed between some variants. These results strongly support the hypothesis that intracellular phagemid abundance serves as the limiting factor for phage secretion, particularly during log phase when cellular resources are most abundant and replication and secretion machinery are most active. Among the sender variants studied, +gp3_p15Aφ, +gp3_pColAφ, and M13KE senders displayed the highest per cell phage secretion rates. Unlike M13KE, +gp3_p15Aφ and +gp3_pColAφ also achieved higher growth rates, making them promising candidates for applications requiring efficient phage secretion and intercellular communication.

Copy numbers of phagemids +gp3_p15Aφ, +gp3_pColAφ, and +gp3_pSC101φ change between the log phase and the stationary phase of growth by 49.61, 36.49, and 4.31 copies per genome. In contrast, phagemids +gp3_pUCφ and +gp3_pBR322φ have similar copy numbers across different growth phases. These differences in observed phagemid copy number likely reflect differences in replication and maintenance of these replication origins as well as the burden of phage secretion. Replication origins of +gp3_pUCφ and +gp3_pBR322φ are derivatives of parental origin ColE1, where replication is regulated by an interplay between the ROP protein and the RNA I/RNA II duplex with a negative feedback loop regulating the plasmid copy number [49]. The +gp3_p15Aφ origin is related to the ColE1 origin and yet is compatible with it [50]. The +gp3_pColAφ (pColA) origin is also a known homolog of ColE1 that regulates replication using a similar mechanism [51]. Unlike the other four phagemids, the replication origin of +gp3_pSC101φ (pSC101) uses short DNA repeats called iterons and encodes for the RepA gene for controlling copy number [52]. Bacteriophage M13 carries its own replication origin that depends on several viral proteins, such as gp2 to initiate and gp5 to regulate replication, thus controlling the plasmid copy number [33].

The effect of different secretion rates for different replication origin phagemids was further explored in co-culture experiments designed to effect intercellular communication of phagemid DNA. We found that phagemid-specific per cell secretion rates influenced the extent and kinetics of receiver cell infection, as revealed by both OD-based growth dynamics (Fig. 4c) and CRISPRi-based transfer frequency measurements (Fig. 5c). Notably, in infected-receivers-only selection conditions, we observed a hierarchy in early receiver growth rates (OD_600_ at 5 h), which closely mirrored the rank order of secretion rates determined earlier. Variants like +gp3_p15Aφ and M13KE rapidly initiated receiver infection and subsequent growth, while +gp3_pUCφ, the lowest-secreting variant, exhibited the slowest and weakest intercellular communication. These results confirm that secretion rates determine the effective ‘broadcast efficiency’ of a sender strain in a population-level communication context.

However, final culture densities at the 18 h time-points did not correlate directly with early infection, especially for the highest-producing phagemids. Instead, we observed an inverse relationship between initial sender inoculum and final infected-receiver growth for +gp3_p15Aφ. This likely reflects a resource competition effect, where highly productive senders deplete shared nutrients during early growth and secretion, thereby limiting the growth of infected receivers later. These findings underscore a key trade-off in our phage-mediated intercellular communication systems: a higher initial secretion rate is accompanied by faster resource consumption in the media and may consequently impair long-term infected receiver proliferation. As a result, senders with lower secretion rates may be better suited for such applications, especially where no selection is applied.

Overall, our study demonstrates that modular phagemid architectures provide a tuneable platform for regulating phage-mediated intercellular communication, with copy number serving as a key lever for optimizing secretion and transfer. The phagemid variants we present—spanning a range of replication origins and copy numbers—offer a versatile toolkit for engineering microbial consortia, studying ecological interactions, and deploying programmable delivery of genetic payloads. Future work could explore selective packaging of orthogonal messages, targeting of specific receivers within a population, further optimisation of phage packaging and secretion rates, and burden-mitigating designs to enhance control and performance.

Summarily, this work lays the foundation for scalable and tuneable DNA communication for payload delivery and synthetic biology applications.

## Methods and Materials

### Bacterial strains, growth conditions, and cloning

All bacteria used in this study are *Escherichia coli* strains (Supplementary Info.). They were grown at 37°C in LB media (liquid with shaking at 180 rpm, or solid LB plates with 1.5% w/v agar) supplemented with the appropriate antibiotics at the following concentrations (unless otherwise indicated): kanamycin (kan 30 μg mL−1), ampicillin (amp 100 μg mL−1), gentamycin (gent 10 μg mL−1), tetracycline (tet 10 μg mL−1) and spectinomycin (spc 50 μg mL−1); concentrations were halved when using multiple antibiotics for selection. Strains and antibiotics used as well as the core parental strains are listed in Supplementary Info.

Cloning was performed by Golden Gate Assembly of PCR-amplified DNA fragments using NEB enzymes: Q5 DNA polymerase (#M0492), BsaI-HFv2 (#R3733) and T4 DNA ligase (#M0202M). *E. coli* strains DH5α and TOP10 were used for cloning. All plasmids constructed were verified by Sanger sequencing. Three SEVA series of vectors with kanamycin resistance (pSEVA291, pSEVA261, pSEVA271) were used as starting points for insertion of the M13ps, gp3 and lacZα. A new phagemid with the SEVA architecture was constructed with a ColA replication origin [53], and named pSEVA29A1 in consultation with the SEVA repository. Plasmids used in this study are listed in Supplementary Info.

Phagemids used in growth, PCN, secretion rate, and cell-to-cell communication assays (antibiotic selection) consist of a bacterial origin of replication, *kanR* for antibiotic selection, M13 packaging signal (ps) essential for phagemid ssDNA packaging, *gp3* encoding for minor coat protein essential for phage secretion and *lacZ*α to easily identify turbid plaques during phage enumeration assays. Control phage M13KE (NEB #E8101S) was also modified to include a *kanR* gene. Phagemids used in the i-CRISPRi experiments had, in addition, sgRNA-1 expressed from a constitutive promoter [40]. These phagemids were made in two versions, one with (+gp3) and one without (-gp3) *gp3* gene. The phagemids were co-transformed with the appropriate helper (pSEVA63_HelperΔgIII for-gp3 phagemids and pSEVA63_Helper for +gp3 phagemids [40]) into *E. coli* Top10 cells to make the senders.

### Sender growth, phage and bacterial pellet isolation

Sender strains (Supplementary Info) were streaked on LBA plates (1.5% w/v agar) and grown overnight at 37°C. Single colonies were inoculated in 5 mL LB media with appropriate antibiotics and incubated overnight at 37°C, with 180 rpm shaking. Overnight cultures were diluted 1000× in 100 mL fresh LB media with antibiotics and incubated for ∼15 h at 37°C, 180 rpm. Periodically, optical density was recorded (OD_600_; spectrophotometer UVisco V-1100D) and 1 mL culture sample was spun down at 4500 ×g for 10 min, supernatant was filtered (0.22 μm filter, Millex SLGP033RS), and the resulting phage preps stored at 4°C and pellets were stored at-80°C. Phage titres at each time-point were estimated using PFU assays (see below) and pellets were used for qPCR analysis (see below)

### Phage counting

Receiver cells (ER2738F_HΔgIII) grown overnight were diluted 1000× and re-grown at 37°C in LB (+tet + gent) until they reached a spectrophotometer OD_600_ between 1 and 1.5. Cells were chilled on ice for at least 30 min and then 90 μL aliquoted into eppendorf tubes. The tubes were moved to RT shortly before mixing 10 μL of different phage dilutions (10^−1^–10^−14^) with the receiver cells and then adding the mix to 10 mL of soft LBA (0.75% w/v agar with 0.2 mM IPTG and 40 μg mL^−1^ X-gal), previously aliquoted into a 15 mL tube and kept molten at 50°C. The phage + receiver mix in the soft agar was immediately poured onto a solid plate with 20 mL hard LBA (1.5% w/v agar), and after the soft LBA had solidified the plate was incubated at 37°C for 16–24 h. Plaques of the non-lytic M13 phage are turbid/ diffused, usually making them harder to see. IPTG and X-gal colours the plaques blue (LacZω in the F-plasmid is complemented by the LacZα in the phagemid), making them easier to visualise. Plaque counts from plates were used to determine the mL^−1^ titres of the phage preps according to the formula: PFU count / (phage dilution * phage volume used in mL).

### Plasmid copy number estimation

Pellet samples stored in 2 mL Eppendorf tubes at-80°C were thawed by adding an appropriate amount of pre-warmed (25°C) elution buffer (EB; NEB #T1010). EB volume to add was calculated to make the theoretical optical density (OD_600_) of the pellet re-suspension 0.1, using the formula: [EBvol.] = [pellet OD_600_] / 0.1. Tubes were vortexed for 5s before incubating them at 95°C for 10 mins. Samples were kept on ice for 1 min before centrifuging at 19,000 xg for 5 mins at room temperature. Supernatant was used for qPCR reactions.

The qPCR reaction was set up to estimate the DNA concentrations of *E. coli* genomic DNA and phagemid DNA by ∼100 bp regions amplified from *gapA* and *gp3* genes with primers listed in Supplementary Info. Standard curves were obtained by using concatenated gapA-gp3 DNA of different predetermined amounts (10-fold dilutions between 0.2 and 0.00002 nM) for qPCR experiments. qPCR amplification was done using Q5 DNA polymerase. A reaction mix of 10 μL comprised 0.2 μM forward (FAM-BBQ650 dual-labelled probe) and reverse primers, 1x Q5 High-Fidelity 2X Master Mix (NEB #M0492), and 2 μL of processed pellet samples. Each reaction was repeated 3 times. Thermocycler conditions were as follows: 3 min initial denaturation at 98 °C and then 40 cycles of 10 s denaturation at 98 °C followed by 30 s of annealing at 69.5 °C and 15 s of extension at 72 °C. The 5’-quencer is hydrolysed by Q5 polymerase before the primer extension, separating the fluorophore from the quencher to emit a signal [54] captured by the qPCR machine (Eppendorf Realplex2 Mastercycler). Raw data is in Supplementary Figs.

### Instantaneous growth and secretion rate analysis

Growth rates between two consecutive time-points were calculated according to the following formula: Specific Growth Rate (μ) = ln (OD2/OD1) / (t2-t1), where OD1 and OD2 are the OD_600_ values at time-points t1 and t2.

Secretion rates between two consecutive time-points were calculated according to the following formula: Secretion Rate = μ * (P2-P1) / (C2-C1), where P1 and P2 are the phage concentrations and C1 and C2 are the cell concentrations at time-points t1 and t2. μ is the specific growth rate calculated above. OD_600_ of sender cells was converted to cell concentration values using the fit based on experiments reported before: CFU = 293192345 * (OD_600_ ^ 1.67) [40].

### Phage-mediated sender-to-receiver communication in co-cultures

For the communication plate-reader experiments, an overnight culture of receiver cells (ER2738F) was diluted 1000× and re-grown to a spectrophotometer OD_600_ of 0.4, following which it was cooled on ice for ∼30 min, pellets were washed, and several OD_600_ dilutions made (0.136, 0.068, 0.034 and 0.0) while still on ice. Overnight culture of senders was diluted 500x and re-grown to a spectrophotometer OD_600_ of 0.2, following which it was cooled on ice for ∼30 min, pellets were washed, and several OD_600_ dilutions made (0.125, 0.062, 0.031, 0.015, 0.007 and 0.0) while still on ice. 90 μL of receiver cell dilutions were added per well to a 96-well plate in quadruplet (for the four different growth conditions), followed by 90 μL of the sender cell dilutions also in quadruplet. The plate was run at 37 °C for 1 h, following which 20 μL of LB with the appropriate antibiotics (10× concentrated, to achieve the 1× end-concentration) was added to each well, and the plate was grown overnight at 37°C, 205 cpm, while recording OD_600_ at 15 min intervals.

### Receiver infection CRISPRi time lapse in growing conditions

Receiver strain (carrying F-plasmid + dCas9-GFP plasmid) and sender strains (carrying helper plasmid + sgRNA phagemid) were streaked on LBA plates (1.5% w/v agar), with appropriate antibiotics, and grown overnight at 37°C. Single colonies were inoculated in 5 mL LB media with appropriate antibiotics and incubated overnight at 37°C, with 180 rpm shaking. Overnight cultures were diluted 1000× in 100 mL fresh LB media with antibiotics and incubated at 37°C, 180 rpm, until they reached OD_600_ ∼0.3. Cultures were cooled on ice for at least 30 min, pellets were washed, and several dilutions were made with different ODs (0.12, 0.06, 0.03 and 0.0), while still on ice. 100 μL of receiver cultures were mixed with 100 μL of sender cultures in a 96-well plate and grown in plate reader for 5 h with no antibiotics while recording OD_600_ and GFP fluorescence at 15 min intervals. At 0, 1, 2, 3, and 4 hours, 10 μL co-culture was added to 1× phosphate-buffered saline (PBS) containing 2 mg mL^−1^ kanamycin to stop protein expression and kill the cells. Following this, the cells in 1× PBS + kan (2 mg mL^−1^) were stored at 4°C for later flow cytometry analysis.

### Phage transfer frequency calculations

Transfer frequencies were calculated from the co-culturing flow cytometry data in Fig. 5. The events recorded at each time-point were gated by fluorescence to obtain the number of cells of each type: senders (S, no GFP), uninfected receivers (Ru, GFP ON) and infected receivers (Ri, GFP OFF). The transfer frequency was calculated as Ri/S*(Ri + Ru), according to the formula used to obtain transconjugant frequency in conjugation experiments [27].

### Flow cytometry analysis

Flow cytometry samples containing cells in 1× PBS (with 2 mg mL^−1^ of kanamycin) were analysed using the Attune NxT flow cytometer (Thermofisher) equipped with a Cytkick autosampler. Samples were fed into the flow cytometer using 96-well plates. Around 20,000 bacterial events were recorded per sample, excluding dead cells and debris from the analysis using FSC and SSC thresholds of 100. GFP fluorescence was measured using excitation by a 488 nm laser and a 530/30 nm filter (BL1 channel). The BL1 fluorescence threshold for each ON/OFF circuit was defined as the lower extreme of the fluorescence distribution from the ON receiver population. The autofluorescent sender population was defined as the upper extreme of the fluorescence distribution from a sender alone population. Voltages used were FSC: 319, SSC: 282, BL1: 292, for all experiments. Data collected were analysed using Attune Cytometric v5.3, and plotted using Python scripts.

